# First record of *Temnothorax convexus* (Forel, 1894) in Portugal (Hymenoptera: Formicidae) with an updated checklist of ants from the country

**DOI:** 10.1101/2021.07.25.453702

**Authors:** Javier Arcos, Diogo Chaves, Paco Alarcón, Ángel Rosado

## Abstract

The arboreal ant *Temnothorax convexus* (Forel, 1894) is reported for the first time in Portugal. A single colony was found nesting inside an abandoned gall near Lisbon. A full morphometric evaluation of the specimens is provided, including a small sample from the recently discovered population in South Iberia. High-definition photographs of the worker caste are given, together with a comparison with the most similar species in Iberia and an update of the Portuguese ant checklist.

## Introduction

The ant species richness in Iberia is among the largest in the Mediterranean basin, including a high rate of endemic taxa (Borowiec, 2014; Janicki *et al*., 2016; Guénard *et al*., 2017). However, only few species are ubiquitous in the whole region, with a great number of taxa being only present in one of the six recognized refugia (Tinaut & Ruano, 2021). In this sense, targeting undersampled refugia appears to be vital in order to produce a final checklist of the Iberian myrmecofauna. The Atlantic refugia, which includes Portugal, is an example of a poorly known region (Borowiec & Salata, 2017), as the continuous finding of previously unrecorded species suggests. The most recent critical checklist of the ant fauna of Portugal listed a total of 106 species (Salgueiro, 2002), which is far from being close to the more than 300 species currently known to inhabit Iberia (Sánchez-García et al., *in prep*.).

During a myrmecological survey conducted by the second author on the outskirts of Lisbon, in the small village of Lousa (municipality of Loures), a colony of a blackish *Temnothorax* species was recovered from an abandoned gall of *Quercus faginea* Lam., containing the queen and approximately 30 workers, together with some female pupae and recently emerged individuals. After a thorough analysis of the workers, we identified the sample as *Temnothorax convexus* (Forel, 1894), representing the first record of this arboreal species in Portugal and the second population known to occurr in Iberia. This finding constitutes also the most northern and western location of the species.

## Material and methods

Measurements were taken using a Nikon SMZ-U stereomicroscope with a magnification between 70 and 225x depending on the character. A cross-scaled ocular micrometre with 100 divisions was used, with an estimated error of 0.01 mm. Specimens were dried and temporarily mounted before measuring and later preserved in alcohol. To minimize measuring errors, characters present on each side of the specimen were both measured and the arithmetic mean was then calculated; when percentual differences greater than 5% were detected for a specific character pair in the same individual this was remeasured to assess whether discordant values represented an occasional recording error or an asymmetry. True asymmetric characters are here defined as characters with a mean percentual asymmetry equal or greater than 3% for all of the worker specimens in the dataset. When any structure is absent, for example an scape, only one side is considered. The following characters were identified as true asymmetric, requiring the evaluation of both sides of the specimens for a reliable interpretation: EL: (4.35% ± 4.15%); LMH (9.19% ± 8.73%); MGr (7.50% ± 23.72%); SPST (6.42% ± 7.62%); PEL (3.21% ± 4.72%); PPL (4.77% ± 3.90%). The variation of the rest of the measured characters is: SL (1.17% ± 1.76%); L1F (0.95% ± 1.80%); ML (1.74% ± 1.32%); MH (1.46% ± 1.32%); PEH (1.02% ± 1.60%); SPEPH (1.58% ± 3.37%); PECW (1.66% ± 2.59%); PPH (1.95% ± 2.97%. Measured characters and indexes are those usually used in the genera *Temnothorax* (Seifert 2006; Csősz et al. 2015), but some modifications and new characters have been introduced. For each individual, 25 characters were recorded (41 primary measurements considering bilaterally present characters). All measurements are in millimeters and follow the format “mean ± standard error (lowest measurement – highest measurement)”. The photographic material consisted of a Sony A6000 camera equipped with a Laowa 25mm ultra-macro lens with 2.5 to 5X magnifications, a Plan APO 10X and Nikon Mplan 20X ELWD microscope objectives. For the focus stacking technique, an electronic macro rail with a resolution < 1 μm per photo was used. Lighting was provided by two 1W LEDs of adjustable intensity.

### Measurements

— HL: maximum cephalic length in the median line, measured in frontal view. The cephalic capsule must be carefully tilted to the position with the true maximum. Excavations of the posterior margin of the head and/or clypeus reduce HL.
— HWb: maximum cephalic width just posterior to the eyes, measured in frontal view.
— CS: cephalic size. The arithmetic means of HL and HWb [(HL/HWb)/2].
— PoOC: postocular distance, measured in frontal view. Using a cross-scaled reticle, position the head as in HL and measure PoOC using as reference (1) the perpendicular projection of the posterior margin of the eye to the median line of head and (2) the median point of the posterior head margin.
— EL: maximum length of the eye, measured in lateral view. Not pigmented ommatidia should also be measured.
— FRS: distance of the frontal carinae immediately caudal of the posterior intersection points between frontal carinae and the lamellae dorsal to the torulus, measured in frontal view (Fig. 1A). If these dorsal lamellae do not laterally surpass the frontal carinae, the deepest point of scape corner pits may be taken as a reference line. These pits take up the inner corner of the scape base when the scape is fully switched caudad and produce a dark triangular shadow in the lateral frontal lobes immediately posterior to the dorsal lamellae of the scape joint capsule.
— SL: maximum straight-line scape length excluding the articular condyle, measured in frontal view.
— L1F: length of distal funiculus segment, measured in frontal view (Fig. 1B). Note that the segment may be caudally or ventrally curved; in this case, it should be measured in lateral view.
— ML: mesosoma length, measured in lateral view. Take as reference points (1) the transition point between anterior pronotal slope and anterior propodeal shield and (2) the caudalmost point of the propodeal lobe, which is usually rounded. In gynes: length from caudalmost point of the propodeal lobe to the most distant point of the steep anterior pronotal face.
— MH: mesosoma height, measured in lateral view. Use a cross-scaled reticle and measure the height using as reference (1) the lowest point of katepistemus and (2) the perpendicular projection to a line connecting the dorsal most points of promesonotum and propodeum.
— MGr: maximum depth of metanotal groove, measured perpendicularly to the tangent connecting the dorsalmost points of promesonotum and propodeum, given as per cent ratio of CS (%) [(MGr/CS)*100].
— PW: maximum pronotal width, measured in dorsal view. In queens, width is measured immediately frontal of the tegulae.
— UHM: count of unilateral hairs on mesosoma excluding those on propodeal spines. To achieve a correct count, the mesosoma must be positioned in fronto-dorsal view to correctly establish its median longitudinal axis. Appressed and/or laterally positioned hairs should also be counted. Not measured in queens, since they are very hairy and UMH in this case is not efficient to measure.
— LMH: length of longest-standing hair on pronotum measured in lateral view.
— SPEPH: maximum subpetiolar process height, measured in lateral view. Take as reference (1) the point where the anterior corner of the subpetiolar process meets the ventral petiole margin and (2) the lowermost point of the structure.
— PEL: petiolar length, measured in lateral view. Use a cross-scaled reticle and take as reference (1) the point where the anterior corner of the subpetiolar process meets the ventral petiole margin and (2) the lower and caudalmost point of the caudal cylinder (Fig. 1D).
— PEH: petiole height, measured in lateral view. Use a cross-scaled reticle to measure the maximum height of the perpendicular projection of a line that extends from (1) the point where the anterior corner of the subpetiolar process meets the ventral petiole margin and (2) the lowermost point of the caudal cylinder.
— PPL: postpetiole length, measured in lateral view. The maximum distance of the postpetiole is measured from the anterior margin to the posterior margin, perpendicular to the dorsal profile line of the postpetiole.
— PPH: maximum height of postpetiole, measured in lateral view (Fig. 1D). It is the longest distance from the deepest point of the subpetiolar process to the superior profile of postpetiole.
— PECW: petiolar cylinder width, measured in lateral view. Use a cross-scaled reticle and take as reference (1) the uppermost point of the petiolar cylinder and (2) the perpendicular projection of the lowermost point of the petiolar cylinder.
— SPBA: the smallest distance between the lateral margins of the spines at their base; it should be measured in dorsofrontal view so that the wider parts of the ventral propodeum do not interfere with the measurement in this position (Fig. 1C); if the lateral margins of spines diverge continuously from the tip to the base, the smallest distance at the base is not defined; in this case, SPBA is measured at the level of the bottom of the interspinal meniscus.
— SPST: distance between the center of propodeal stigma and spine tip; the stigma center refers to the midpoint defined by the outer cuticular ring but not to the center of the real stigma opening that may be positioned eccentrically.
— SPWI: maximum distance between the outer margins of the propodeal spines, measured in dorsal view.
— PEW: maximum petiole width, measured in dorsal view.
— PPW: maximum postpetiole width, measured in dorsal view.

**Figure 1.**
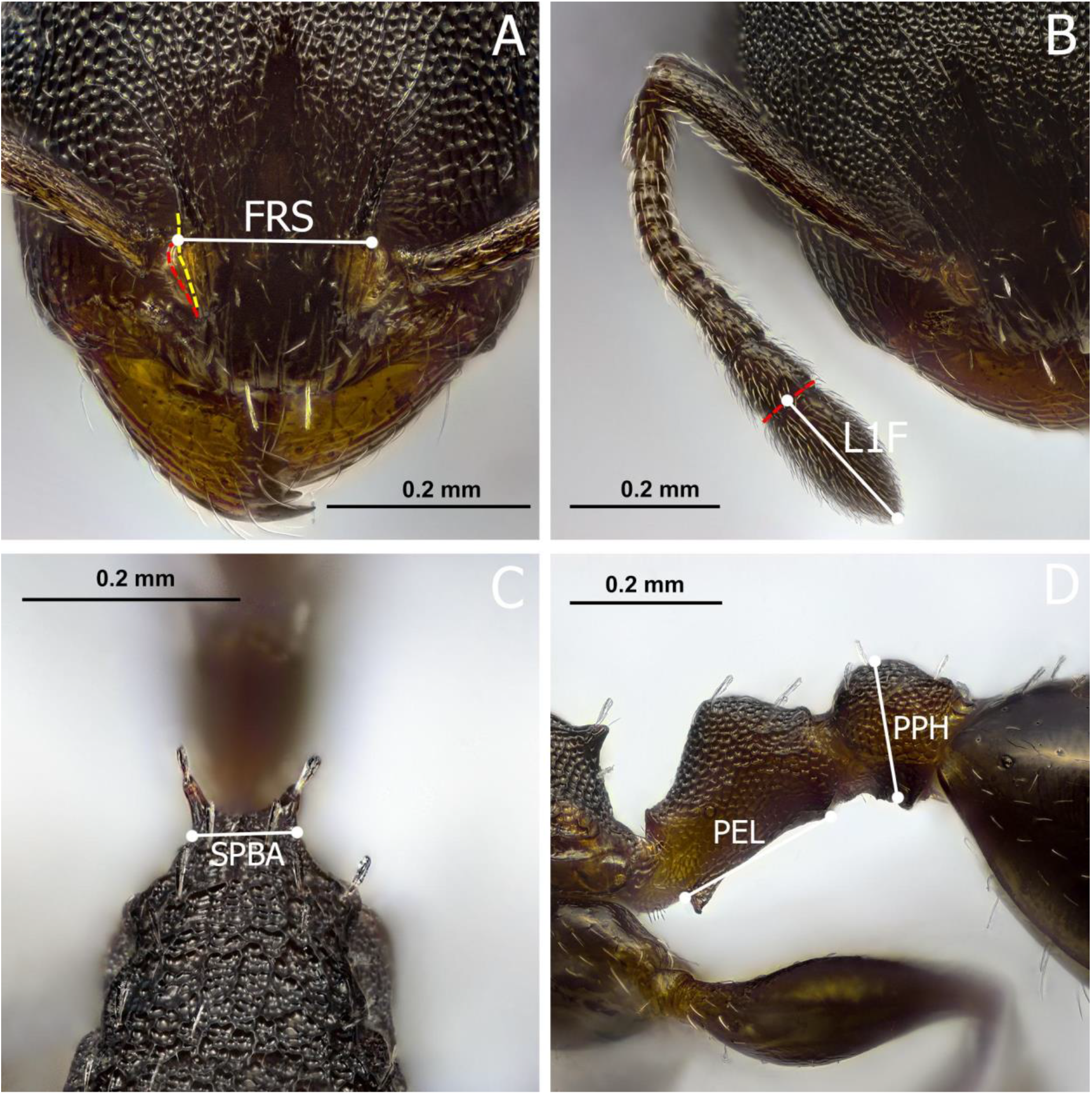
Worker of *T. convexus*. A: frontal view showing how to measure FRS. B: frontal view showing how L1F is measured. C: detailed view of the mesosoma in dorso-frontal view, the position in which SPBA is measured. D: lateral view of the petiole and postpetiole, with the measuring points for PEL and PPH.

## Results

A single colony was found nesting in an abandoned gall of *Quercus faginea* Lam., containing the queen and approximately 30 workers, together with some female pupae that were reared in the laboratory. The investigated sample was identified as *T. convexus* according to the original description, the morphologic description provided by Guillem & Bensusan (2019) and by comparisson with the imaged syntype of the species in Antweb.org (CASENT0909022). *T. convexus* belongs to the *angustulus* group according to the literature (Cagniant & Espadaler, 1997; Galkowski & Cagniant, 2017). This is mainly integrated by arboreal species with the following character combination: (1) petiole shape triangular in lateral view, (2) pronounced sculpture on mesosoma consisting on ground reticulation with superimposed longitudinal striae and (3) dark coloration of mesosoma with occasional reddish pronotum in some bicoloured species. However, one of the main diagnostic features of the group (1) does not fit the characteristics found in *T. convexus*, since the petiole in this species is low and rounded. Species-groups in Temnothorax are informally used and few have been properly characterized, and so we are not here proposing a new redefinition of the *T. angustulus* group as we understand that a wider revision of this species is needed in order to determine its diagnostic features.

### Worker diagnosis

#### Investigated material

seven workers from Lousa (Portugal), 38°52’5’.2”N 9°12’48.6”W, ?.V.2021, 230m, Diogo leg., nest in gall; 3 workers from El Puerto de Santa María (Cádiz, Spain), 36°34’17.4”N 6°12’59.7”W, 2m, Rafael Obregon leg., nest in fallen branch. Measurements in millimeters (n = 10): HL: 0.635 ± 0.031 (0.570–0.693) mm; HWb: 0.499 ± 0.032 (0.453–0.564) mm; PoOC: 0.246 ± 0.015 (0.222–0.268) mm; EL: 0.126 ± 0.007 (0.111–0.139) mm; FRS: 0.192 ± 0.013 (0.176–0.213) mm; SL: 0.472 ± 0.029 (0.425–0.527) mm; L1F: 0.222 ± 0.008 (0.208–0.236) mm; LMH: 0.062 ± 0.009 (0.048–0.083) mm; ML: 0.714 ± 0.039 (0.656–0.786) mm; MH: 0.292 ± 0.024 (0.254–0.333) mm; MGr: 0.001 ± 0.002 (0–0.007) mm; SPST: 0.174 ± 0.014 (0.157–0.194) mm; SPL: 0.112 ± 0.015 (0.083–0.139) mm; PEL: 0.236 ± 0.025 (0.208–0.287) mm; PEH: 0.204 ± 0.015 (0.179–0.231) mm; NOL: 0.161 ± 0.014 (0.139–0.185) mm; SPEPH: 0.038 ± 0.006 (0.028–0.046) mm; PECW: 0.130 ± 0.008 (0.120–0.139) mm; PPL: 0.141 ± 0.010 (0.129–0.157) mm; PPH: 0.168 ± 0.012 (0.152–0.194) mm; USH: 9.385 ± 1.293 (6–10.5) mm; SPTI: 0.180 ± 0.019 (0.153–0.213) mm; SPWI: 0.191 ± 0.018 (0.159–0.224) mm; SPBA: 0.132 ± 0.019 (0.110–0.166) mm; PW: 0.341 ± 0.022 (0.313–0.383) mm; PEW: 0.139 ± 0.013 (0.116–0.166) mm; PPW: 0.188 ± 0.014 (0.162–0.213) mm. See Table 1 for the indexes.

**Table 1.** Indexes of *T. convexus* workers and queens.

|  | *T. convexus* from Portugal (n = 7) | *T. convexus* from Cádiz (n = 3) | All worker specimens (n = 10) | Queens from Portugal (n = 2) |
| --- | --- | --- | --- | --- |
| CS | 0.638 ± 0.056<br>[0.577, 0.742] | 0.626 ± 0.025<br>[0.598, 0.644] | 0.634 ± 0.048<br>[0.577, 0.742] | 0.695 ± 0.013<br>[0.686, 0.704] |
| CL/CWb | 1.130 ± 0.031<br>[1.078, 1.172] | 1.145 ± 0.061<br>[1.075, 1.186] | 1.135 ± 0.039<br>[1.075, 1.186] | 1.041 ± 0.019<br>[1.027, 1.054] |
| PoOC/CL | 0.377 ± 0.010<br>[0.364, 0.392] | 0.361 ± 0.012<br>[0.351, 0.375] | 0.373 ± 0.013<br>[0.351, 0.392] | 0.374 ± 0.003<br>[0.372, 0.377] |
| FRS/CS | 0.318 ± 0.013<br>[0.304, 0.337] | 0.341 ± 0.010<br>[0.331, 0.350] | 0.325 ± 0.016<br>[0.304, 0.350] | 0.333 ± 0.006<br>[0.329, 0.338] |
| SL/CS | 0.690 ± 0.008<br>[0.681, 0.702] | 0.704 ± 0.015<br>[0.691, 0.721] | 0.694 ± 0.012<br>[0.681, 0.721] | 0.660 ± 0.031<br>[0.638, 0.682] |
| EL/CS | 0.245 ± 0.013<br>[0.233, 0.269] | 0.241 ± 0.010<br>[0.233, 0.252] | 0.244 ± 0.012<br>[0.233, 0.269] | 0.283 ± 0.001<br>[0.283, 0.284] |
| L1F/CS | $0.338 \pm 0.021$<br>[0.294, 0.361] | $0.336 \pm 0.007$<br>[0.331, 0.341] | $0.337 \pm 0.019$<br>[0.294, 0.361] | $0.287 \pm 0.024$<br>[0.270, 0.304] |
| LMH /CS | $0.081 \pm 0.006$<br>[0.076, 0.091] | - | $0.081 \pm 0.006$<br>[0.076, 0.091] | $0.113 \pm 0.002$<br>[0.112, 0.115] |
| ML/CS | $1.269 \pm 0.031$<br>[1.221, 1.325] | $1.264 \pm 0.014$<br>[1.255, 1.281] | $1.267 \pm 0.027$<br>[1.221, 1.325] | $1.264 \pm 0.035$<br>[1.559, 1.649] |
| MH/CS | $0.554 \pm 0.009$<br>[0.540, 0.563] | $0.544 \pm 0.015$<br>[0.527, 0.558] | $0.551 \pm 0.011$<br>[0.527, 0.563] | $0.917 \pm 0.060$<br>[0.875, 0.959] |
| MGr/CS | $0.149 \pm 0.124$<br>[0.070, 0.402] | $0.000 \pm 0.000$<br>[0.000, 0.000] | $0.104 \pm 0.125$<br>[0.070, 0.402] | - |
| SPST/CS | $0.202 \pm 0.009$<br>[0.193, 0.219] | $0.212 \pm 0.010$<br>[0.201, 0.219] | $0.205 \pm 0.010$<br>[0.193, 0.219] | $0.200 \pm 0.004$<br>[0.197, 0.203] |
| PEL/CS | $0.408 \pm 0.015$<br>[0.389, 0.426] | $0.407 \pm 0.027$<br>[0.388, 0.438] | $0.408 \pm 0.018$<br>[0.388, 0.438] | $0.447 \pm 0.018$<br>[0.434, 0.459] |
| PEH/CS | $0.338 \pm 0.009$<br>[0.327, 0.351] | $0.340 \pm 0.018$<br>[0.326, 0.360] | $0.339 \pm 0.011$<br>[0.326, 0.360] | $0.330 \pm 0.011$<br>[0.322, 0.338] |
| SPEPH/CS | $0.043 \pm 0.005$<br>[0.035, 0.048] | $0.040 \pm 0.008$<br>[0.032, 0.047] | $0.042 \pm 0.006$<br>[0.032, 0.048] | $0.067 \pm 0.001$<br>[0.066, 0.068] |
| PECW/CS | $0.248 \pm 0.010$<br>[0.233, 0.263] | $0.249 \pm 0.009$<br>[0.241, 0.259] | $0.248 \pm 0.009$<br>[0.233, 0.263] | $0.267 \pm 0.005$<br>[0.263, 0.270] |
| PEH/PECW | $1.365 \pm 0.033$<br>[1.323, 1.407] | $1.365 \pm 0.046$<br>[1.313, 1.394] | $1.365 \pm 0.034$<br>[1.313, 1.407] | $1.238 \pm 0.018$<br>[1.225, 1.250] |
| PEW/CS | $0.272 \pm 0.026$<br>[0.248, 0.325] | $0.261 \pm 0.013$<br>[0.248, 0.273] | $0.269 \pm 0.022$<br>[0.248, 0.325] | $0.287 \pm 0.004$<br>[0.284, 0.289] |
| PPL/CS | $0.241 \pm 0.004$<br>[0.232, 0.244] | $0.257 \pm 0.006$<br>[0.252, 0.264] | $0.246 \pm 0.009$<br>[0.232, 0.264] | $0.294 \pm 0.024$<br>[0.276, 0.311] |
| PPH/CS | $0.319 \pm 0.017$<br>[0.304, 0.350] | $0.318 \pm 0.022$<br>[0.295, 0.338] | $0.319 \pm 0.017$<br>[0.295, 0.350] | $0.357 \pm 0.007$<br>[0.351, 0.362] |
| PPW/CS | $0.363 \pm 0.030$<br>[0.335, 0.425] | $0.365 \pm 0.009$<br>[0.357, 0.374] | $0.364 \pm 0.025$<br>[0.335, 0.425] | $0.413 \pm 0.008$<br>[0.408, 0.419] |
| SPWI/CS | $0.191 \pm 0.022$<br>[0.171, 0.238] | $0.178 \pm 0.009$<br>[0.171, 0.187] | $0.187 \pm 0.020$<br>[0.171, 0.238] | $0.300 \pm 0.004$<br>[0.297, 0.303] |
| PW/CS | $0.623 \pm 0.027$<br>[0.589, 0.675] | $0.622 \pm 0.023$<br>[0.605, 0.647] | $0.622 \pm 0.024$<br>[0.589, 0.675] | $0.933 \pm 0.018$<br>[0.921, 0.946] |
| USH/CS | $18.435 \pm 3.054$<br>[15.142, 0.223] | - | $18.435 \pm 3.054$<br>[15.142, 24.948] | - |
| SPBA | $0.185 \pm 0.017$<br>[0.171, 0.223] | $0.165 \pm 0.013$<br>[0.155, 0.180] | $0.179 \pm 0.018$<br>[0.155, 0.223] | $0.300 \pm 0.004$<br>[0.297, 0.303] |

Big and robust species (mean CS 0.634 mm); polymorphism more pronounced than other Palearctic species, with a notable size difference between smaller and bigger specimens. Slightly bicolored species with brownish mesosoma and darker head, with slightly darkened antennal clubs and femora. Scapes relatively short (mean SL/CS 0.694). Eyes moderately large (mean EL/CS 0.244). Standing hairs on mesosoma relatively short (mean LMH/CS 0.081). Promesonotal groove distinct in dorsal view, especially in the bigger specimens (Fig. 2D). Metanotal groove absent to very shallow (mean MGr/CS 0.104%). Mesosoma profile concave, with a pronounced decline of the dorsal profile line near the propodeal spines (see Fig. 2A). Propodeal spines strongly reduced to triangular teeth (mean SPST/CS 0.205) and relatively close in dorsal view (mean SPWI/CS 0.187) (see Fig. 1C). Subpetiolar process short but distinct (SPEPH/CS 0.042). Petiole long (mean PEL/CS 0.408) and low (mean PEH/CS 0.339), with rounded apex; some specimens may show a slightly truncated apex. Sculpture of head consisting of reticulate ground sculpture with shallow superimposed longitudinal striae, with absent or very narrow smooth median region on frons. Mesosoma with reticulate ground sculpture and strong longitudinal striae, especially pronounced on sides of pronotum and dorsum. When comparing the measured individuals from Lisbon and Cádiz, mean indexes are concordant. The morphologic coincidence of these two samples with the imaged syntype of *T. convexus* from Forêt de Msila (Algeria) (CASENT0909022) is also complete. Note that a noticeable difference in propodeal spine length is found between both sides of an important number of workers (Fig. 1A), which highlights the necessity of measuring both spines in this species.

**Figure 2.**
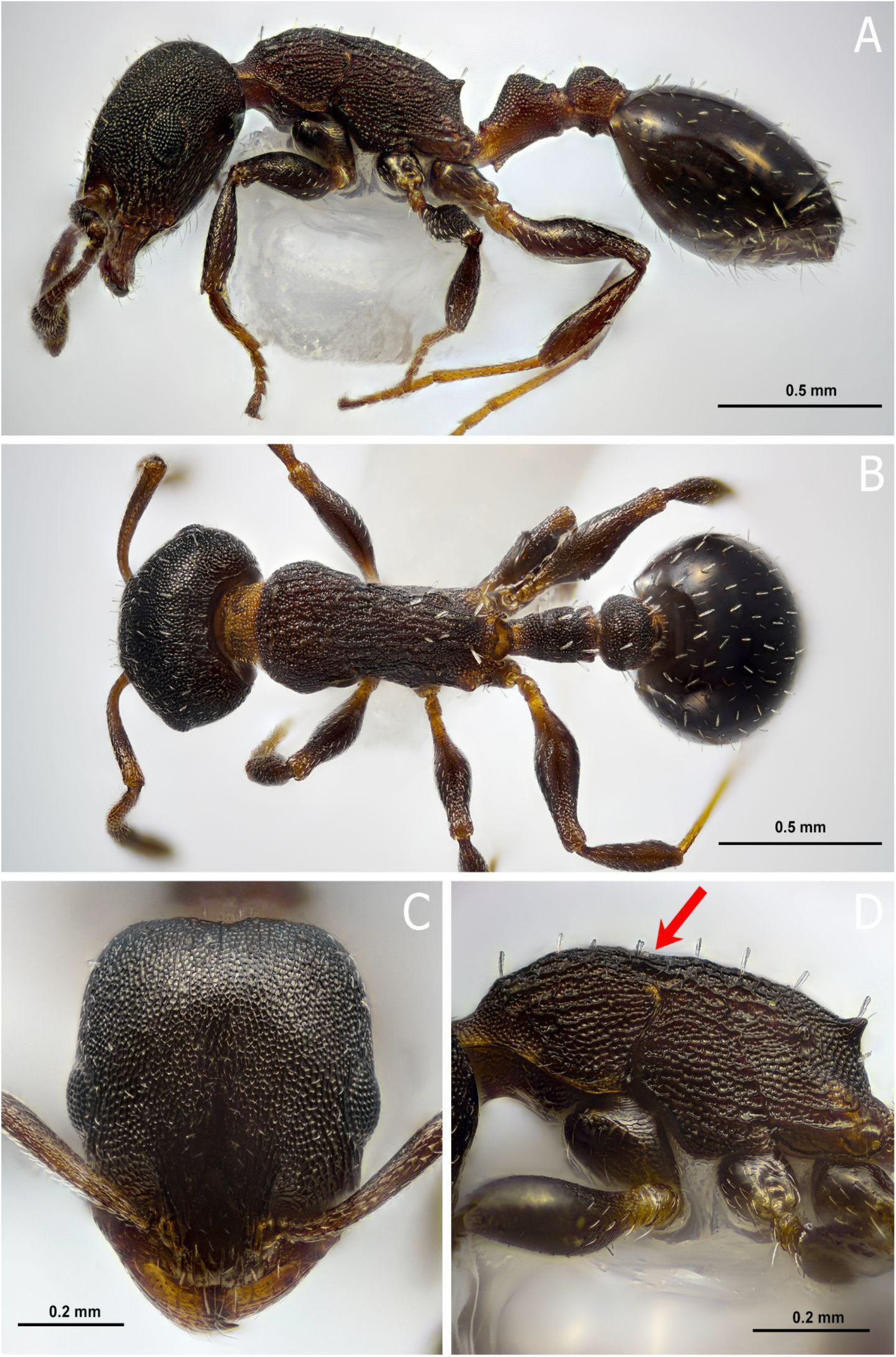
Worker of *T. convexus*. A: lateral view. B: dorsal view. C: frontal view. D: detail of mesosoma, with the distinct short pilosity and pronounced promesotonal furrow (red arrow).

### Queen diagnosis

Two specimens from Lisbon were available for measuring (same collection data as workers) and indexes are presented in Table 1. Measurements (n = 2): 0.695 ± 0.013; HL 0.709 ± 0.020 (0.695, 0.723) mm; HWb 0.681 ± 0.007 (0.677, 0.686) mm; PoOC 0.265 ± 0.005 (0.262, 0.269) mm; FRS 0.232 ± 0.000 (0.232, 0.232) mm; SL 0.459 ± 0.013 (0.450, 0.468) mm; EL 0.197 ± 0.003 (0.195, 0.199) mm; L1F 0.199 ± 0.013 (0.190, 0.209) mm; LMH 0.079 ± 0.000 (0.079, 0.079) mm; ML 1.129 ± 0.003 (1.126, 1.131) mm; MH 0.637 ± 0.029 (0.616, 0.658) mm; SPST 0.139 ± 0.000 (0.139, 0.139) mm; PEL 0.311 ± 0.007 (0.306, 0.315) mm; PEH 0.229 ± 0.003 (0.227, 0.232) mm; SPEH 0.046 ± 0.000 (0.046, 0.046) mm; PECW 0.185 ± 0.000 (0.185, 0.185) mm; PPL 0.204 ± 0.013 (0.195, 0.213) mm; PPH 0.248 ± 0.010 (0.241, 0.255) mm; SPWI 0.209 ± 0.007 (0.204, 0.213) mm; SPBA 0.209 ± 0.007 (0.204, 0.213) mm; PW 0.649 ± 0.000 (0.649, 0.649) mm; PEW 0.199 ± 0.007 (0.195, 0.204) mm; PPW 0.287 ± 0.000 (0.287, 0.287) mm.

Queens are visually bigger than other Palearctic species of the genus (mean CS 0.120 mm). Head dorsum brownish with slightly darkened antennal clubs and femora; mesosoma brownish with yellowish areas. Note that queens can be both uniformly dark or bicolored according to Guillem & Bensusan (2019). Our specimens have yellowish pronotum and the rest of the mesosoma brownish. Mesosoma curved, with convex scutum and scutellum in lateral view. Scape relatively short (mean SL/CS 0.660). Standing hairs on mesosoma short (mean LMH/CS 0.113). Mesosoma profile slightly concave. Propodeal spines strongly reduced to two obtuse angles, without distinct denticles. Subpetiolar process short but distinct (SPEPH/CS 0.067). Petiole very long (mean PEL/CS 0.447) and low (mean PEH/CS 0.330), with rounded apex. Sculpture of head consisting of reticulate ground sculpture with shallow superimposed longitudinal striae. Mesosoma with reticulate ground sculpture and strong longitudinal striae, especially pronounced on sides of pronotum and scutum.

## Discussion

*T. convexus* was originally described from Algeria (Forel, 1894). Emery (1915) described *Leptothorax submuticus* from Tangier (Morocco), which was later synonymized with *T. convexus* (Cagniant & Espadaler, 1997). Recently, Guillem & Bensusan (2019) recorded it from several locations in Gibraltar and Cádiz (South Iberia). In its revision of the *Temnothorax* from Morocco, Cagniant & Espadaler (1997) couldn1t find any *T. convexus* sample in the region and raised the possibility of the species representing a teratogenic form of *Temnothorax atlantis* (Santschi, 1911). According to Galkowski & Cagniant (2017), *T. convexus* has not been located again in the type locality. The population from Cap Espartel (Morocco), described as *Leptothorax convexus var. timida* (Santschi, 1912), was synonymized under *T. algiricus* (Forel, 1894) by Cagniant & Espadaler (1997); however, the imaged worker holotype (CASENT0912920) strongly recalls the general appearance of *T. convexus* and does not match the concept of *T. algiricus*, a taxon with long propodeal spines and triangular petiole, instead of short propodeal spines and rounded petiolar node as seen in the mentioned type.

In Iberia, its closest relatives based on the morphology of the worker and queen castes are *Temnothorax angustulus* (Nylander, 1856), *Temnothorax aveli* (Bondroit, 1918), *Temnothjorax continentalis* Galkowski & Cagniant, 2017, *Temnothorax corticalis* (Schenck, 1852) and *Temnothorax nadigi* (Kutter, 1925). These are mostly arboreal species that have coarse sculpture on the mesosoma and short to medium length propodeal spines. A comparison between all five species and *T. convexus* is presented in Table 2. In general terms, the length of the propodeal spines, which are extremely reduced in *T. convexus*, could exclude *T. angustulus* and *T. continentalis* from the differential diagnosis of the worker caste, since they are usually longer and curved; the petiolar shape is also very different in *T. convexus*. The workers of the species *T. aveli* and *T. corticalis* are of yellowish mesosoma, thus differentiating them easily from the dark brown mesosoma of *T. convexus*. It should be noted that the presence of the species *T. corticalis* in Iberia is dubious, since it has not been found in the region in the last 30 years (since De Haro & Collingwood, 1991), which raises the possibility of a misidentification with *T. aveli* or other closely-related arboreal species, especially since some records are based on the queen caste only (Collingwood & Yarrow, 1969), which is very similar in both taxa. Interestingly, *T. convexus* and *T. nadigi* are two apparently unrelated species that share some uncommon traits within the Iberian members of the genus *Temnothorax*, including greatly reduced propodeal spines, coarse striation on mesosoma and low petiole, and to some extent the characteristic convex mesosoma. However, they are easily separated based again on the yellowish aspect of the mesosoma of *T. nadigi* and its long-standing hairs on the mesosoma, compared to the very short hairs of *T. convexus*.

**Image 1.**
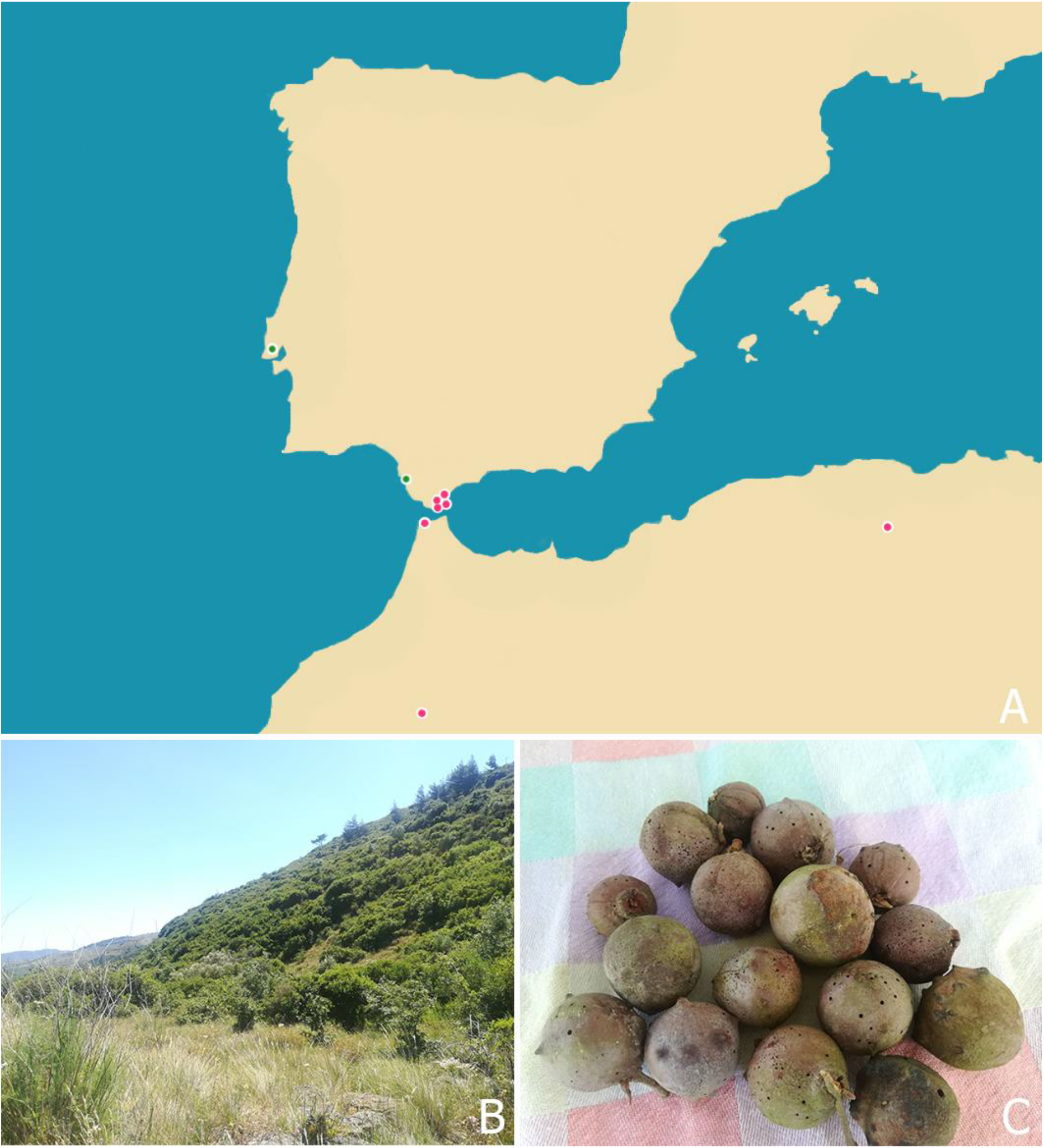
A: known records of *T. convexus* (pink circles) and new locations (green circle). B: general view of the habitat where the sample from Lisbon was found. C: galls from the same place where the sample was located.

**Table 2.** Morphological differences between the worker caste of *T. convexus* and its most similar species in Iberia.

|  |  |  |
| --- | --- | --- |
| *Temnothorax convexus*<br>Head entirely reticulated, with superficial superimposed longitudinal striae on frons. Propodeal spines reduced to short denticles. Petiolar apex rounded. Metanotal groove absent to shallow. Mesosoma convex in lateral view. Bicolored with lighter mesosoma and slightly reddish pronotum. | vs. | *T. angustulus* and *T. continentalis*<br>Posterior half of the head mostly smooth and shiny, with superimposed longitudinal striae on frons. Propodeal spines triangular and distinct. Petiolar apex distinctly triangular. Metanotal groove usually distinct in profile view. Mesosoma flat in lateral view. Concolorous mesosoma in the case of *T. angustulus* and markedly reddish in *T. continentalis*. |
| *T. convexus*<br>Petiole apex rounded. Hairs on mesosoma short. Workers of dark coloration and lighter brownish mesosoma. | vs. | *T. aveli*<br>Petiole apex triangular. Hairs on mesosoma long. Workers usually bicolored, with a dark brown head and yellowish mesosoma. |
| *T. convexus*<br>Petiole in lateral view long and rounded. Species of dark brown coloration, with brownish mesosoma and occasional reddish pronotum. | vs. | *T. corticalis*<br>Petiole in lateral view short and triangular. Bicolored species with yellowish mesosoma and darker head. |
| *T. convexus*<br>Antennal club not markedly darkened. Sculpture of head uniform, frons included. Hairs on mesosoma short. Bicolored with brownish mesosoma and darker head and gaster, occasionally with reddish pronotum. Petiole apex usually rounded. | vs. | *T. nadigi*<br>Antennal club clearly darkened. Sculpture of frons reduced, with a smooth and shiny medium strip. Hairs on mesosoma longer. Bicolored with yellowish mesosoma and darker head. Petiole apex usually truncated. |

The queen caste of *T. convexus* is also very distinct. Queens of *T. angustulus* and *T. continentalis* have well developed propodeal spines, of triangular aspect and wide base, and the shape of the petiole is high and triangular, contrasting with the almost absent propodeal spines and low and rounded petiolar node of T. convexus. On the other hand, queens of *T. aveli*, *T. convexus*, *T. corticalis* and *T. nadigi* have very reduced propodeal spines. From *T. aveli*, the queen of *T. convexus* is distinguished by the overall darker coloration (usually light brown mesosoma in *T. aveli*) and the reduction of the propodeal spines to two very obtuse angles (small denticles in *T. aveli*). The queen of *T. convexus* is easily separated from *T. corticalis* based on the low and long petiole of the first in contrast with the high and short triangular petiole of the later (Seifert, 2018). The most similar queen is found in the species *T. nadigi*, but the pilosity is significantly longer and the antennal clubs are darkened in the case of *T. nadigi*, while they are not distinctly darkened in *T. convexus*. The curved mesosoma is also found to some extent in the queen caste of *T. convexus*, which is a very prominent feature within the Iberian *Temnothorax* queens.

We suspect that the record of *T. atlantis* (Santschi, 1911) from Portugal that appears in Henin *et al*. (2000) could actually be *T. convexus*. We base this assumption on the following facts: (1) the locality were *T. atlantis* was found is at some 60 km from our new record of *T. convexus*, (2) the species *T. atlantis* is similar to *T. convexus* based on the morphology of the worker caste and a misidentification with the latter is plausible, especially when *T. convexus* was apparently not considered in the differential diagnosis by these authors, (3) the authors of the paper state that the shape of the petiole is not coincident with the original description of *T. atlantis* since it is “more concave”, a feature that matches the petiole shape of *T. convexus*, (4) the fact that *T. atlantis* is a North African species and its dubious presence in Iberia is solely based on the Portuguese record and (5) there is no other Iberian species similar to *T. atlantis* apart from *T. convexus* and so the possibility of a third species being involved in the differential diagnosis is very unlikely. A visit by the second author (3.VI.21) to the area where *T. atlantis* was originally recorded did not provide any sample of *Temnothorax*. On the contrary, the area was found to be infested by *Linepithema humile* (Mayr, 1868), an invasive exotic species which could have in fact eradicated any *Temnothorax* population inhabiting the locality. However, more effort is needed to assess this possibility, as *T. convexus* appears to be an uncommon species even in the areas where it is present (Guillem & Bensusan, 2019).

Since the checklist published by Salgueiro (2002), who recognized 106 species of ants in continental Portugal, the discovery of new species in the country has not stopped. The addition of *T. convexus* to the Portuguese ant fauna follows this trend. The species that have been located in the country in the last decades are summarized in Table 3 in chronological order, raising the final number of continental Portuguese ants to 136. Some new nomenclatural changes affecting the Portuguese fauna are worth mentioning here. The recently described *Temnothorax alfacarensis* (Tinaut & Reyes-López, 2020) was already recorded as *Temnothorax sp*. in Espadaler *et al*. (2008). Previous records of *Temnothorax specularis* (Emery, 1916) should now be treated as *Temnothorax exilis* (Emery, 1869) according to Salata *et al*. (2018). The identity of the species integrating the *Tapinoma nigerrimum* species-complex should be carefully studied, as the old records of *T. nigerrimum* (Nylander, 1856) could actually represent more than one species and not necessarily the true *T. nigerrimum* (see Seifert *et al*., 2017). The following species are noted from Portugal in Borowiec & Salata (2012) but the original reference or record has not been found in the present paper and are therefore not counted in our updated checklist: *Aphaenogaster subterranea* (Latreille, 1798), *Aphaenogaster testaceopilosa* (Lucas, 1849), *Cataglyphis cursor* (Fonscolombe, 1846), *Cryptopone ochracea* (Mayr, 1855), *Lasius neglectus* Van Loon, Boomsma & Andrasfalvy, 1990, *Liometopum microcephalum* (Panzer, 1798), *Pheidole indica* Mayr, 1879 and *Tapinoma subboreale* Seifert, 2012.

**Table 3.** Summarized additions to the continental Portuguese ant fauna in the last decades since Salgueiro (2002), in chronological order. The last column represents the accumulative number of species.

| Paper | Additions | Species |
| --- | --- | --- |
| Salgueiro (2002) | *Stenamma debile* (Forster, 1850), *Lasius piliferus* (Seifert, 1992), *Myrmica wesmaeli* (Bondroit, 1918) | 109 |
| Boeiro *et al.* (2002) | *Stigmatomma gaetulicum* (Baroni Urbani, 1978), *Goniomma kugleri* (Espadaler, 1986), *Oxyopomyrmex saulcyi* (Emery, 1889), *Hypoponera abeillei* (André, 1881) | 113 |
| Peira *et al.* (2002) | *Temnothorax angustulus* | 114 |
| Salgueiro (2003) | *Temnothorax formosus* (Santschi, 1909), *Tetramorium impurum* (Forster, 1850). *Lasius lasioides* (previously omitted in Salgueiro, 2002). | 117 |
| Seifert (2003) | *Cardiocondyla nigra* (Forel, 1905) | 118 |
| Espadaler *et al.* (2008) | *Bothriomyrmex saundersi* (Santschi, 1922), *Leptanilla revelierii* (Emery, 1870), *Temnothorax parvulus* (Schenck, 1852), *Temnothorax alfacarensis* (Tinaut & Reyes-López, 2020) | 122 |
| Boeiro *et al.* (2009) | *Strumigenys silvestrii* (Emery, 1906), *Temnothorax albipennis* (Curtis, 1854), *Lasius fuliginosus* (Latreille, 1798) | 125 |
| Obregón & Reyes (2012) | *Nylanderia jaegerskioeldi* (Mayr, 1904) | 126 |
| Branstetter (2012) | *Stenamma sardoum* (Emery, 1915) | 127 |
| Espadaler & Gómez (2014) | *Tetramorium biskrense* (Forel, 1904) | 128 |
| Gonçalves et al. (2014) | *Strongylognathus caeciliae* (Forel, 1897), *Temnothorax tyndalei* (Forel, 1909) | 130 |
| Zina & Franco (2015) | *Plagiolepis xene* (Stärcke, 1936) | 131 |
| Garcia *et al.* (2015) | *Stenamma westwoodii* (Westwood, 1839) | 132 |
| Borowiec & Salata (2017) | *Goniomma baeticum* (Reyes & Rodriguez, 1987), *Myrmoxenus kraussei* (Emery, 1915), *Temnothorax luteus* (Forel, 1874) | 135 |
| Seifert *et al.* (2017) | *Tapinoma ibericum* (Santschi, 1925) | 136 |
| Present work | *Temnothorax convexus* | 137 |

## Acknowledgements

Special thanks to Joaquín López Reyes for the loaned specimens of *T. convexus* from Cádiz.

## Notes

### Competing Interest Statement

The authors have declared no competing interest.

